# Interoceptive sensitivity modulates heart rate and insula activity when listening to music

**DOI:** 10.1101/2021.12.05.471331

**Authors:** Toru Maekawa, Takafumi Sasaoka, Toshio Inui, Shigeto Yamawaki

**Affiliations:** Center for Brain, Mind and KANSEI Sciences Research, Hiroshima University, 1-2-3 Kasumi, Minami-Ku, Hiroshima, 734-8551, Japan; Otemon Gakuin University, 2-1-15, Nishi-ai, Ibaraki, Osaka 567-8502, Japan

**Keywords:** Emotions, Heart rate, Insula activity, Interoceptive sensitivity, Music

## Abstract

Interoception plays an important role in emotion processing, but the relationship between the physiological responses associated with emotional experience and interoception is unclear. In this study, we measured interoceptive sensitivity using the heartbeat discrimination task and investigated the effects of individual differences in interoceptive sensitivity on changes in heart rate and insula activity in response to music-induced emotions. We found that the heart rate increased when listening to the music pieces rated as emotionally high-touching in the high interoceptive sensitivity group only. Compared to the emotionally low-touching music, listening to the emotionally high-touching music was associated with higher insula activity. Furthermore, relative to individuals with low interoceptive sensitivity, the region of interest analysis of the insula subregions for individuals with high interoceptive sensitivity revealed significant activity in the bilateral dorsal granular insula, the right ventral dysgranular insula, and the right granular and dorsal dysgranular insula while listening to the high-touching music pieces. Our results suggest that individuals with high interoceptive sensitivity use their physical condition to assess their emotional level when listening to music. Furthermore, the insula activity may reflect the use of interoception to estimate emotions.

## 1. Introduction

Interoception refers to the sense of the internal state of the body (Craig, 2002; Ceunen *et al.*, 2016) and has attracted attention due to its important role in emotions, decision making, and social interaction (Pollatos *et al.*, 2005; Craig, 2008; Seth, 2013; Critchley and Garfinkel, 2017; Khalsa *et al.*, 2018). One of the objective ways to investigate the relationship between emotions and interoception is to examine physiological responses when experiencing emotions. Pollatos *et al.* (2007) and Pollatos and Schandry (2008) reported a significant decrease in the heart rate after the presentation of emotional images to participants with high interoceptive sensitivity. Herbert *et al.* (2010) also showed that the heart rate and cardiovascular parameters during emotional image observation differ depending on the level of interoceptive sensitivity. These results indicate that the physiological response to emotional images changes according to interoceptive sensitivity. In contrast, a previous study reported no differences in heart rate and SCR in response to the presentation of emotional videos between the high and low interoceptive sensitivity groups (Wiens *et al.*, 2000). Therefore, the relationship between emotional and physiological responses and interoceptive sensitivity is not always consistent.

In the study of interoception, heartbeats are widely used to examine individual differences in interoceptive sensitivity. The most popular method is the heartbeat counting task (Schandry, 1981), in which participants count their heartbeat in mind within a certain period. The previous studies of emotion and physiological response used the heartbeat counting task to estimate the interoceptive sensitivity. However, this method has been criticized as the participants’ knowledge of heart rate affects their scores (Murphy *et al.*, 2018; Zamariola *et al.*, 2018). To address these issues, some studies have used the heartbeat discrimination task (Garfinkel *et al.*, 2015; Ring and Brener, 2018) in which participants listen to the sound stimulus and respond whether it matches the timing of their heartbeat. In this task, it is thought that the participants’ knowledge and strategy do not affect the results, and the participants’ sensitivity to the heartbeat can be measured more accurately than with the heartbeat counting task (Ring and Brener, 2018).

Neuroimaging studies have strongly suggested a link between interoception and emotion (Quigley *et al.*, 2021). The insula is considered central to interoceptive processing, and insula activity during interoceptive tasks (Critchley *et al.*, 2004; Simmons *et al.*, 2013) and the relationship between insula activity and individual differences in interoceptive sensitivity (Kuehn *et al.*, 2016; Tan *et al.*, 2018) have been demonstrated. The insula is also considered to be an important area for linking interoception with emotion or social interaction (Gu *et al.*, 2013), and overlapping activations of emotional and interoceptive tasks have been reported (Zaki *et al.*, 2012).

The insula is categorized into three types according to its cytoarchitectural organization: agranular cortex located in the anterior region, dysgranular cortex located in the dorsal-mid region, and granular cortex located in the posterior region (Mesulam and Mufson, 1982). A meta-analysis of insula activity showed that the social-emotional, cognitive, and sensorimotor tasks activated the anterior-ventral, the anterior-dorsal, and the mid-posterior insula, respectively (Kurth *et al.*, 2010). However, consistent results have not been obtained regarding the relationship between individual differences in interoceptive sensitivity and the activities of the insula subregions. Previous studies have reported a correlation between the activity of the right anterior insula and interoceptive sensitivity (Critchley *et al.*, 2004; Haruki and Ogawa, 2021), a correlation between the activity of the mid insula and interoceptive sensitivity (Tan *et al.*, 2018), and decreased network centrality in the right posterior insula in the high interoceptive sensitivity group (Kuehn *et al.*, 2016). These inconsistencies might be due to the difference in experimental tasks. A meta-analysis of the brain activity during various interoception-related tasks reported that the commonly activated regions are the mid and posterior insula (Schulz, 2016). However, the relationship between the physiological responses associated with emotional experience and individual differences in interoception has not been fully clarified.

Based on these previous findings, we made the following hypotheses: (1) the physiological response during the emotional experience of music correlates with individual differences in interoceptive sensitivity; (2) the interoceptive sensitivity measured by the heartbeat discrimination task is related to the physiological response; and (3) the activity of the posterior insula during an emotional experience correlates with individual differences in interoceptive sensitivity. To test these hypotheses, we conducted an experiment in which participants listened to various music pieces while measuring brain activity using functional magnetic resonance imaging (fMRI) and heart rate as a physiological response. Music is often used as a visual stimulus for emotion induction (Gilet, 2008). A comparison of visual and auditory emotional induction showed that emotional recognition of visual stimuli is more accurate, whereas musical stimuli induce larger changes in electrodermal activity (EDA), heart rate, and brain wave amplitude than visual stimuli (Baumgartner *et al.*, 2006). Therefore, it is expected that the relationship between emotions and interoception will be more prominent when music is used to induce emotions.

## 2. Methods

### Participants

The study involved 52 participants (31 females, 21 males, age range: 18–35 years, average age: 22.6 ± 2.81 years). All participants were right-handed, not professional musicians, and had never majored in music. One participant was excluded from the analysis due to a malfunction in the physiological recording. The participants provided written informed consent in accordance with the Declaration of Helsinki and received a monetary reward for participating. The research ethics committee of Hiroshima University approved this study (approval number: E-965).

### Heartbeat counting task

In the heartbeat counting task, the participants counted and reported the number of heartbeats between two beep sounds. The sound intervals comprised 6 conditions of 25, 30, 35, 40, 45, and 50 s, and once for each condition. During the task, a 3-lead electrocardiograph (ECG) was attached to the chest of the participants and the ECG was recorded at 1000 Hz using a Biopac MP160 System (Biopac Systems, Goleta, CA).

From the participants’ responses, the interoceptive accuracy (IA) score was calculated using the following formula:

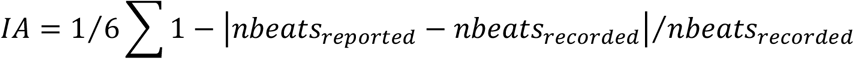

Here, *nbeat_recorded_* represents the correct number of heartbeats, and *nbeat_reported_* represents the number of heartbeats reported by the participant.

### Heartbeat discrimination task

The heartbeat discrimination task is used to determine whether the timing of the beep sound and one’s own heartbeat are synchronized when a beep sound is slightly delayed from the actual heartbeat (Whitehead *et al.*, 1977). A beep sound was presented immediately after the participant’s R-peak detected by the ECG (0 ms condition), or with 150 ms (150 ms condition), 300 ms (300 ms condition), and 450 ms (450 ms condition) delay from the R-peak timing. In each trial, participants listened 10 beep sounds and answered whether the beep timing matched or did not match their heartbeat. Six trials under each condition were performed, totaling 24 trials.

In the heartbeat discrimination task, participants do not always feel that their heartbeats are simultaneous with the sounds when the delay time is 0 ms (Ring and Brener, 2018). In addition, since the heartbeat is periodic, it might be impossible to distinguish whether the stimulus is delayed or advanced in some cases. Therefore, we created a novel heartbeat discrimination index. To characterize the performance of the participants, we approximated the Gaussian function to the simultaneous judgment ratio (Figure 1). In this fitting, the periodicity of the heartbeat was dealt with as follows: In the case of a participant with an R-R interval of 800 ms, as an example, because the heartbeat occurs in an almost constant cycle, the delay of the sound of 450 ms is indistinguishable from the advance of the sound of 350 ms (= 800–450 ms). Therefore, the response data under the 450 ms delay condition can also be treated as relevant data when the sounds are presented 350 ms earlier than the heartbeats. Each participant’s data at five points, (450 – RRI) ms, 0 ms, 150 ms, 300 ms, and 450 ms, were fitted to the following Gaussian function.

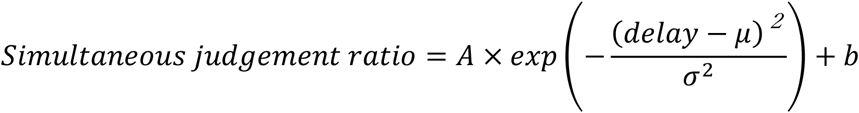

**Figure 1.**
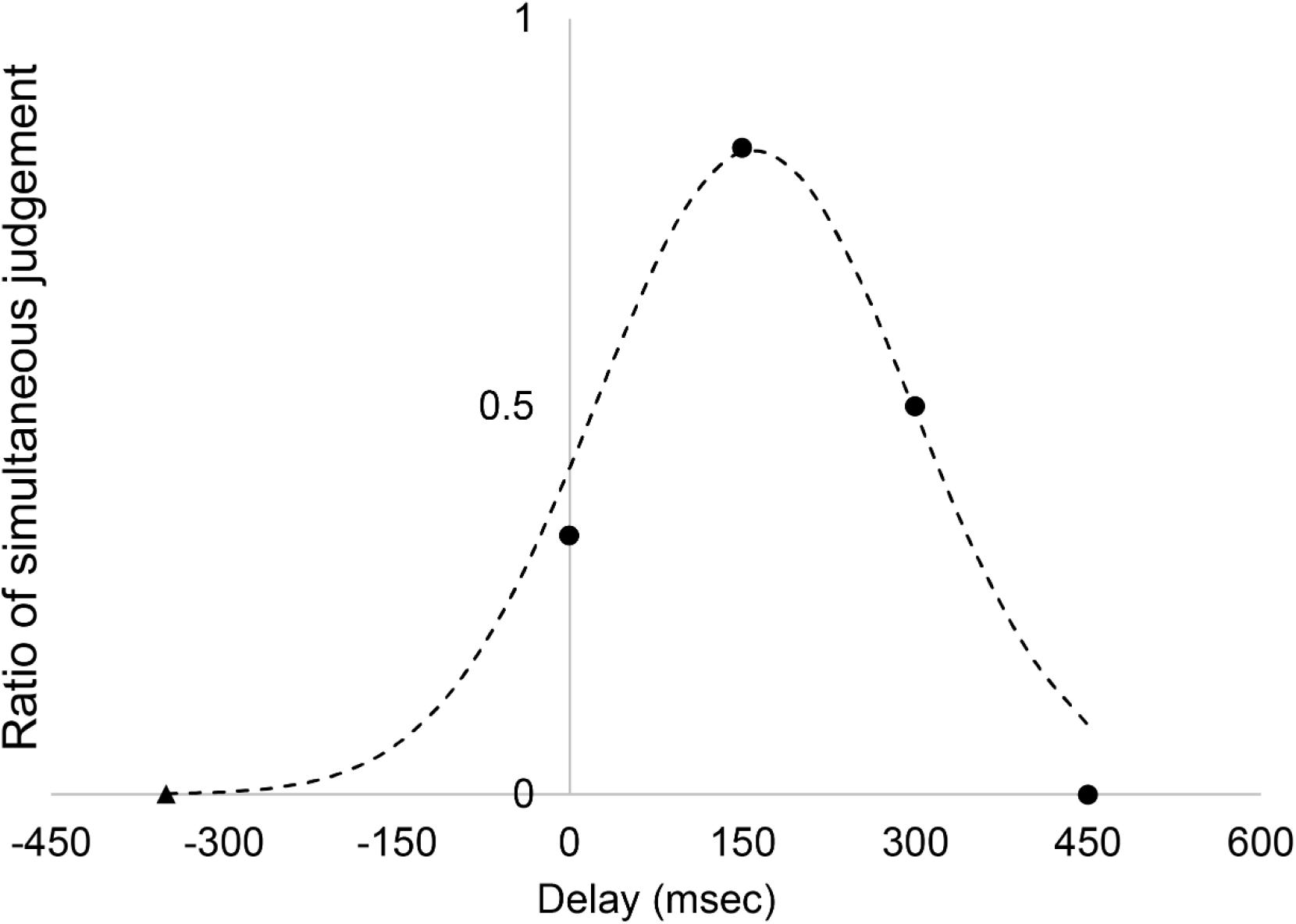
A schematic diagram of the analysis of the heartbeat discrimination task. The horizontal axis shows the delay time of the beep from R-peak, while the vertical axis shows the percentage of trials in which participants responded that the timing was matched. The triangular symbol indicates the extrapolated condition considering the periodicity of the heartbeat (see details in the text). The dotted line shows the Gaussian approximation.

There are four fitting parameters: *A, μ, σ,* and *b.* If a participant has a good heartbeat sensitivity, the Gaussian shape should be higher and sharper. Therefore, we used amplitude *A* and variance *σ* as indicators of interoceptive sensitivity (IS) for subsequent analyses.

### Music task

Participants listened to a music piece for 30 s using an MRI scanner and evaluated emotional touching. Musical stimuli were selected from the music used by Proverbio *et al.* (2015). To widen the range of touching, we selected 10 tonal music pieces as high-emotionally touching music and 10 atonal music pieces as low-emotionally touching music (Table 1). In addition, for the 10 tonal music pieces, two variations were made by changing the pitches to one half note higher and three half notes lower; then, a music piece was created by synthesizing them (Koelsch *et al.*, 2006). Most of these synthesized music pieces were dissonant and unpleasant; thus, the touching level was expected to be low. Therefore, a total of 30 music pieces were used, including 10 tonal, 10 atonal, and 10 pitch-shifted tonal music pieces.

**Table 1.**
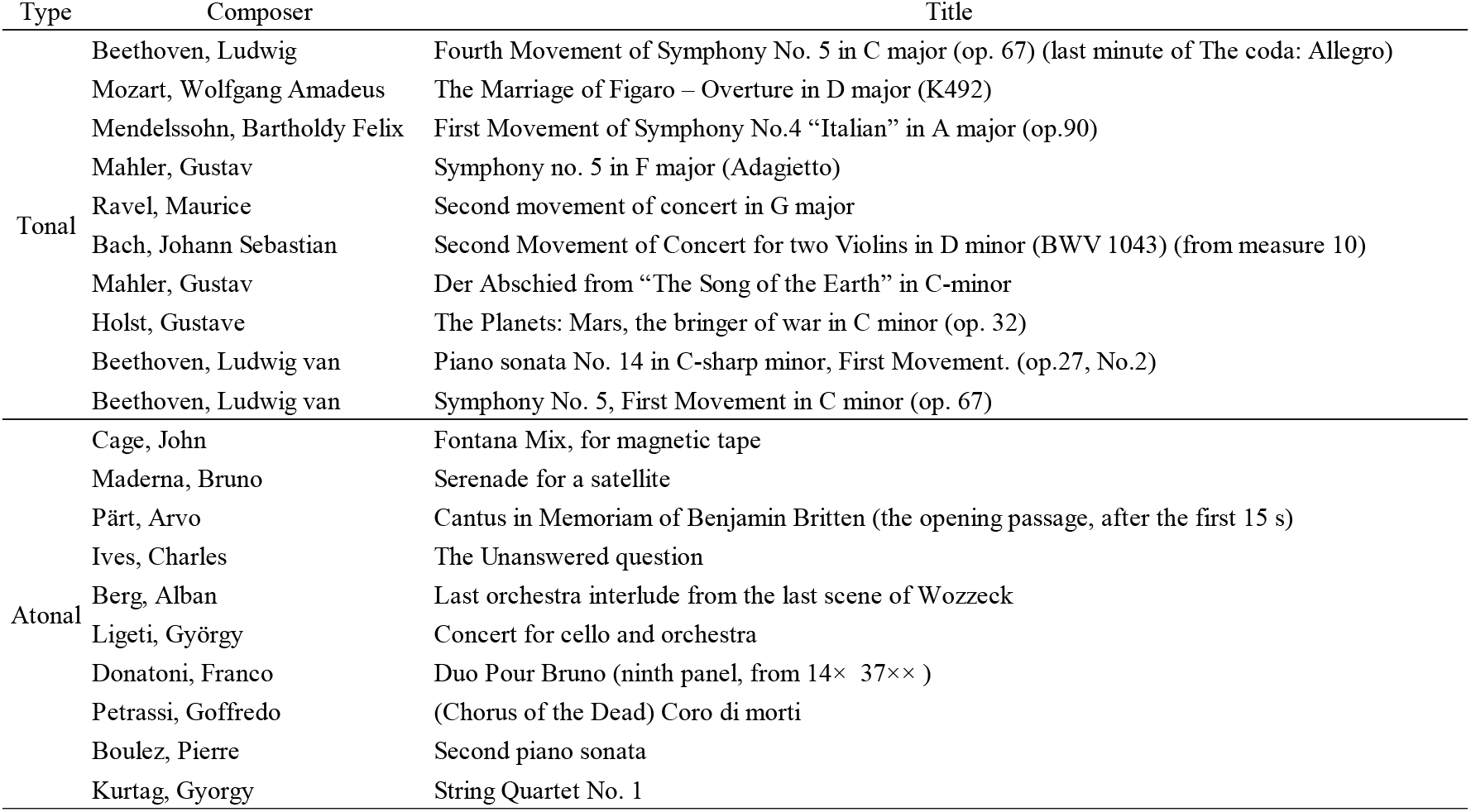
List of music used in the experiment.

| Type | Composer | Title |
| --- | --- | --- |
| Tonal | Beethoven, Ludwig | Fourth Movement of Symphony No. 5 in C major (op. 67) (last minute of The coda: Allegro) |
|  | Mozart, Wolfgang Amadeus | The Marriage of Figaro – Overture in D major (K492) |
|  | Mendelssohn, Bartholdy Felix | First Movement of Symphony No.4 “Italian” in A major (op.90) |
|  | Mahler, Gustav | Symphony no. 5 in F major (Adagietto) |
|  | Ravel, Maurice | Second movement of concert in G major |
|  | Bach, Johann Sebastian | Second Movement of Concert for two Violins in D minor (BWV 1043) (from measure 10) |
|  | Mahler, Gustav | Der Abschied from “The Song of the Earth” in C-minor |
|  | Holst, Gustave | The Planets: Mars, the bringer of war in C minor (op. 32) |
|  | Beethoven, Ludwig van | Piano sonata No. 14 in C-sharp minor, First Movement. (op.27, No.2) |
|  | Beethoven, Ludwig van | Symphony No. 5, First Movement in C minor (op. 67) |
| Atonal | Cage, John | Fontana Mix, for magnetic tape |
|  | Maderna, Bruno | Serenade for a satellite |
|  | Pärt, Arvo | Cantus in Memoriam of Benjamin Britten (the opening passage, after the first 15 s) |
|  | Ives, Charles | The Unanswered question |
|  | Berg, Alban | Last orchestra interlude from the last scene of Wozzeck |
|  | Ligeti, György | Concert for cello and orchestra |
|  | Donatoni, Franco | Duo Pour Bruno (ninth panel, from 14× 37×× ) |
|  | Petrassi, Goffredo | (Chorus of the Dead) Coro di morti |
|  | Boulez, Pierre | Second piano sonata |
|  | Kurtág, György | String Quartet No. 1 |

Participants listened to a music piece while undergoing the fMRI. They wore earplugs and headphones with a noise-canceling function (OptoActive, Optoacoustics Ltd, Or Yehuda, Israel). In addition, a photoplethysmogram was recorded from the participants’ left index finger at 1000 Hz using Biopac MP 150 system. In each trial, after a presentation of a fixation cross for 5 s, participants listened a music piece for 30 s and rated touching level by a visual analog scale. Each music piece was presented once, and participants performed 30 trials. The experiment was divided into two sessions of 15 trials, with a 10-minute break between sessions.

### fMRI data acquisition and analysis

A 20-channel head coil 3T MRI system (Siemens MAGNETOM Skyra, Siemens LTD., Erlangen) was used for data acquisition. Before the experiment, a high-resolution T1-weighted structural scan was obtained using a three-dimensional magnetization prepared rapid gradient echo imaging sequence (TR = 2300 ms, TE = 2.98 ms, flip angle = 9 °, field of view = 256 mm, voxel size 1 × 1 × 1 mm, 176 slices). The functional images were obtained using an echo-planar T2*-weighted multiband gradient echo sequence (TR = 1000 ms, TE = 30 ms, flip angle = 80 degrees, field of view = 192 mm, voxel size 3.0 × 3.0 x 3.2 mm, 42 slices, acceleration factor 3). The experiment was divided into two blocks, and 707–902 functional image data were obtained for each block. The first 10 functional images for each block were excluded from the analysis to allow for T1-equilibration. The movement of the head during the task was corrected, normalized using the Montreal Neurological Institute (MNI) template, and smoothed with a 3D Gaussian filter (full width at half maximum 8 mm).

After preprocessing, the first-level analysis was conducted using a general linear model (GLM) for each participant. The GLM comprised regressors for the periods during the presentation of music, presentation of fixation before music was played, presentation of music rated as high-touching, and presentation of music rated as low-touching score, as well as during a response, including pressing buttons. We then obtained the contrast between high-touching trials and low-touching trials. The statistical significance of the contrast was verified by a *t*-test in the group analysis. The significance level was *p* < 0.001 without correction at the voxel-level and *p* < 0.05 with family wise error correction at the cluster level. Next, we conducted a region of interest (ROI) analysis to investigate the effects of individual differences in interoceptive sensitivity. We defined six subregions of the insula for each hemisphere using the Brainnetome Atlas (Fan *et al.*, 2016). For each subregion, we examined whether there was any difference in activity between the high and low groups of interoceptive sensitivity.

## 3. Results

### Heartbeat tasks

The average IA score for the heartbeat counting task was 0.72 ± 0.22. For the heartbeat discrimination task, Gaussian parameters were estimated for each participant, and the average values for each parameter were as follows: *A*: 0.72 ± 0.16, *μ*: 197 ± 265ms, *σ*: 671 ± 520ms, and *b*: 0.26 ± 0.17. We then calculated the correlation between IA and the heartbeat discrimination task parameters. This revealed a weak positive correlation between the amplitude parameter (*A*) only and IA (*A*: *r* = 0.27, *t* (49) = 1.94, *p* = 0.05, *μ*: *r* = −0.03, *t* (49) = 0.21, *p* = 0.86, *σ*: *r* = −0.12, *t* (49) = 0.82, *p* = 0.41, *b*: *r* = −0.19, *t* (49) = 1.33, *p* = 0.09).

### Heart rate during the music task

First, we examined whether the heart rate during the presentation of the music piece would differ depending on the ratings of touching. For each trial, we calculated the average heart rate during the presentation of the fixation cross before the start of the presentation of a music piece as the baseline, and the difference from the baseline as the heart rate. Then, for each participant, we divided 30 trials into three touching-level groups based on the rated scores of touching: high-touching trials, 10 trials in which the touching score was the highest; low-touching trials, 10 trials in which the touching score was the lowest; and middle-touching trials, 10 trials in which the touching score was between low and high. We then averaged the heart rate during the presentation of the music piece for trials in each group. Figure 2 shows the average heart rate for all participants. A one-way repeated-measures analysis of variance (ANOVA) with a factor of touching level group (high-touching, middle-touching, and low-touching trials) was performed on the average heart rate for 30 s while listening to the music piece. This revealed a significant main effect of touching (*F* (1, 50) = 5.69, *p* = 0.02). The post-hoc test with Bonferroni correction revealed a significantly higher heart rate in the high-touching trials than in the low-touching trials (*p* = 0.05). However, a one-way repeated-measures ANOVA with a factor of music type (tonal, atonal, and discorded) revealed no significant main effect (*F* (1, 50) = 0.45, *p* = 0.83).

**Figure 2.**
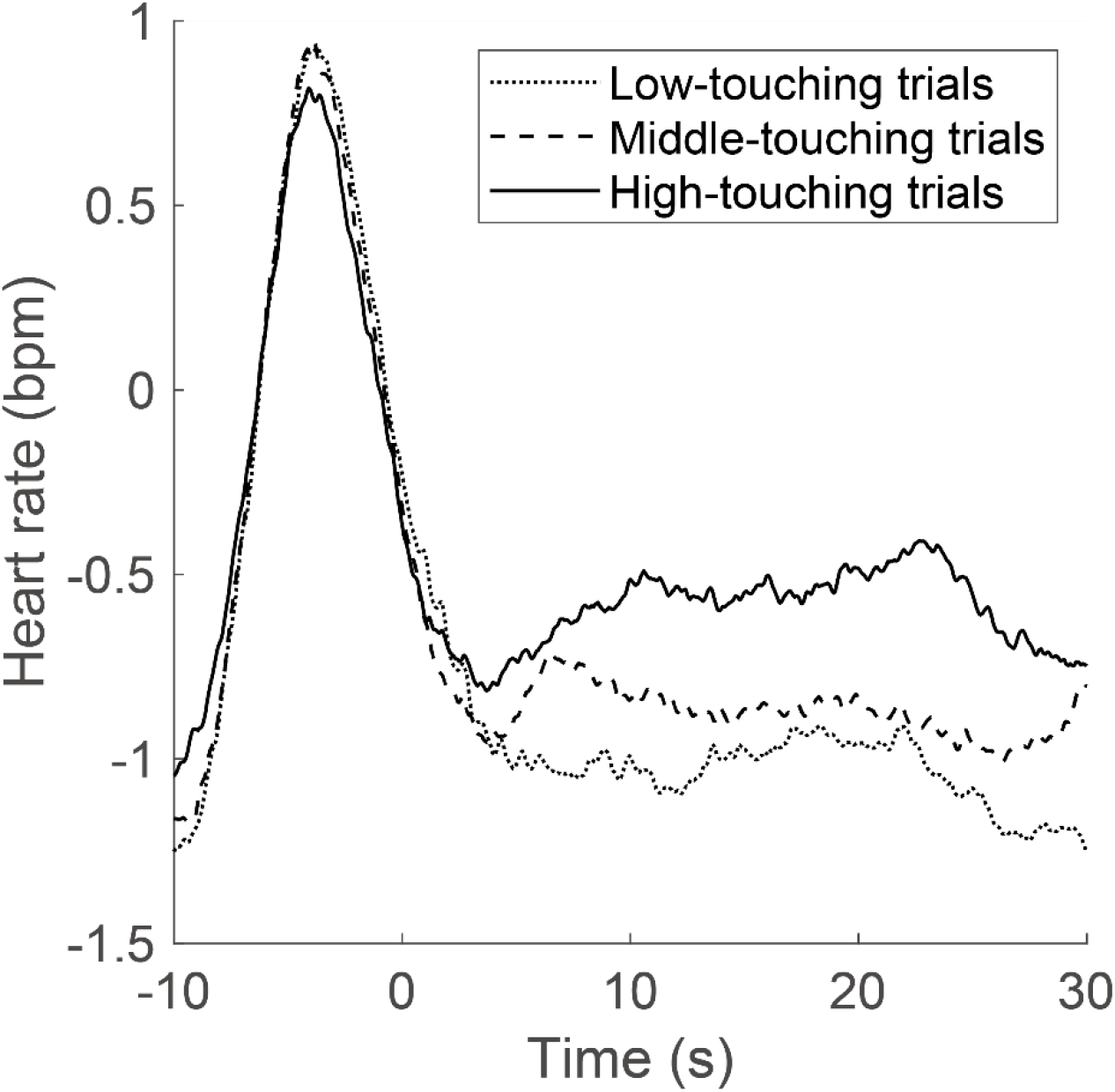
Changes in heart rate when listening to music in the music task. The vertical axis shows the change in heart rate based on the fixation time before the trial. The zero on the horizontal axis indicates the time when the music started.

Next, we divided the participants into two groups according to the heartbeat discrimination task amplitude, that is, fitting parameter *A.* Based on the median of *A,* 25 participants with *A* above the median were classified into the high interoceptive sensitivity (high-IS) group. The remaining 26 participants were classified into the low interoceptive sensitivity (low-IS) group. The average heart rate for each interoceptive sensitivity group is shown in Figure 3. A two-way repeated measures ANOVA revealed a significant interaction between the interoceptive sensitivity group and the touching level group with regard to the average heart rate (IS group: *F* (1, 49) = 0.04, *p* = 0.95; touching level group: *F* (1, 49) = 5.89, *p* = 0.02; IS×touching interaction: *F* (1, 49) = 4.53, *p* = 0.04). The post-hoc test with Bonferroni correction revealed a significant difference between the high-touching trials and the low-touching trials in the high-IS group only (*p* = 0.01).

**Figure 3.**
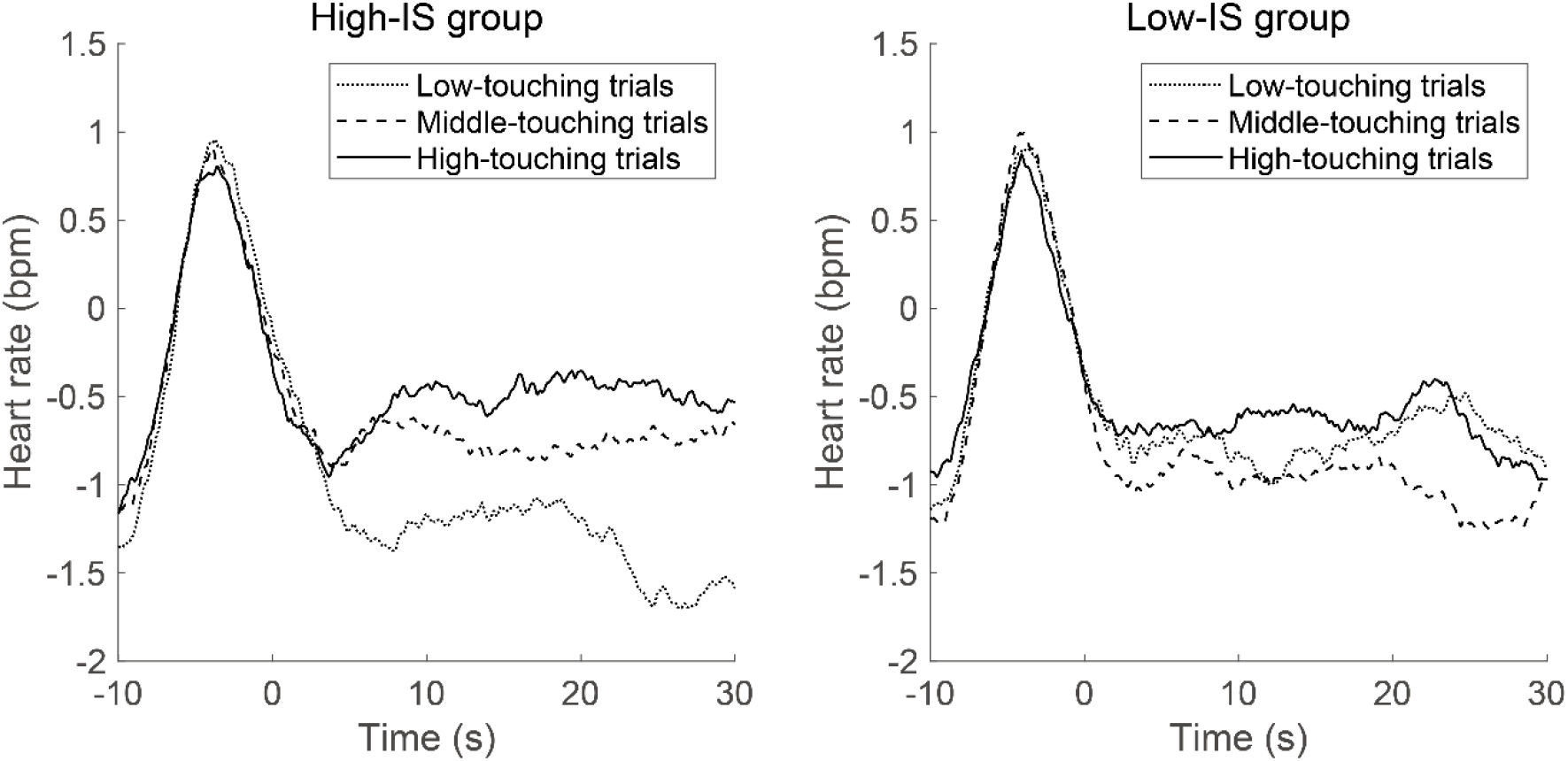
Changes in heart rate grouped according to the interoceptive sensitivity defined by the performance of the heartbeat discrimination task (HDT). Participants were divided into two groups based on the HDT amplitude parameter. The left figure shows the average result of the high-IS group, while the right figure shows the average result of the low-IS group.

We performed the same analyses based on grouping according to the heartbeat counting task score (IS group: *F* (1, 49) = 1.55, *p* = 0.22; touching level group: *F* (1, 49) = 5.56, *p* = 0.02; IS×touching interaction: *F* (1, 49) = 1.07, *p* = 0.31) and the variance parameter (*σ*) of the heartbeat discrimination task (IS group: *F* (1, 49) = 0.18, *p* = 0.67; touching level group: *F* (1, 49) = 5.85, *p* = 0.02; IS×touching interaction: *F* (1, 49) = 1.41, *p* = 0.24), but none of them showed a significant interaction. Therefore, we used the heartbeat discrimination task amplitude parameter (*A*) as an index of interoceptive sensitivity.

Next, we examined whether the touching score was affected by individual differences in interoceptive sensitivity. This analysis revealed a weak but significant positive correlation between interoceptive sensitivity and touching score (*r* = 0.27, *t* (49) = 1.92, *p* = 0.05, Figure 4). In contrast, no significant correlation was found between interoceptive sensitivity and the standard deviation (SD) or the variance of the touching score (SD: *r* = 0.16, *t* (49) = 1.11, *p* = 0.27, variance: *r* = 0.23, *t* (49) = 1.62, *p* = 0.09). Furthermore, to determine whether the heart rate changed due to interoceptive sensitivity regardless of the touching level, we examined the relationship between heart rate changes and interoceptive sensitivity. However, no significant correlation was found between interoceptive sensitivity and average, SD, or variance of the heart rate changes (average: *r* = −0.03, *t* (49) = 0.21, *p* = 0.83; SD: *r* = −0.02, *t* (49) = 0.14, *p* = 0.88; variance: *r* = −0.06, *t* (49) = 0.41, *p* = 0.67).

**Figure 4.**
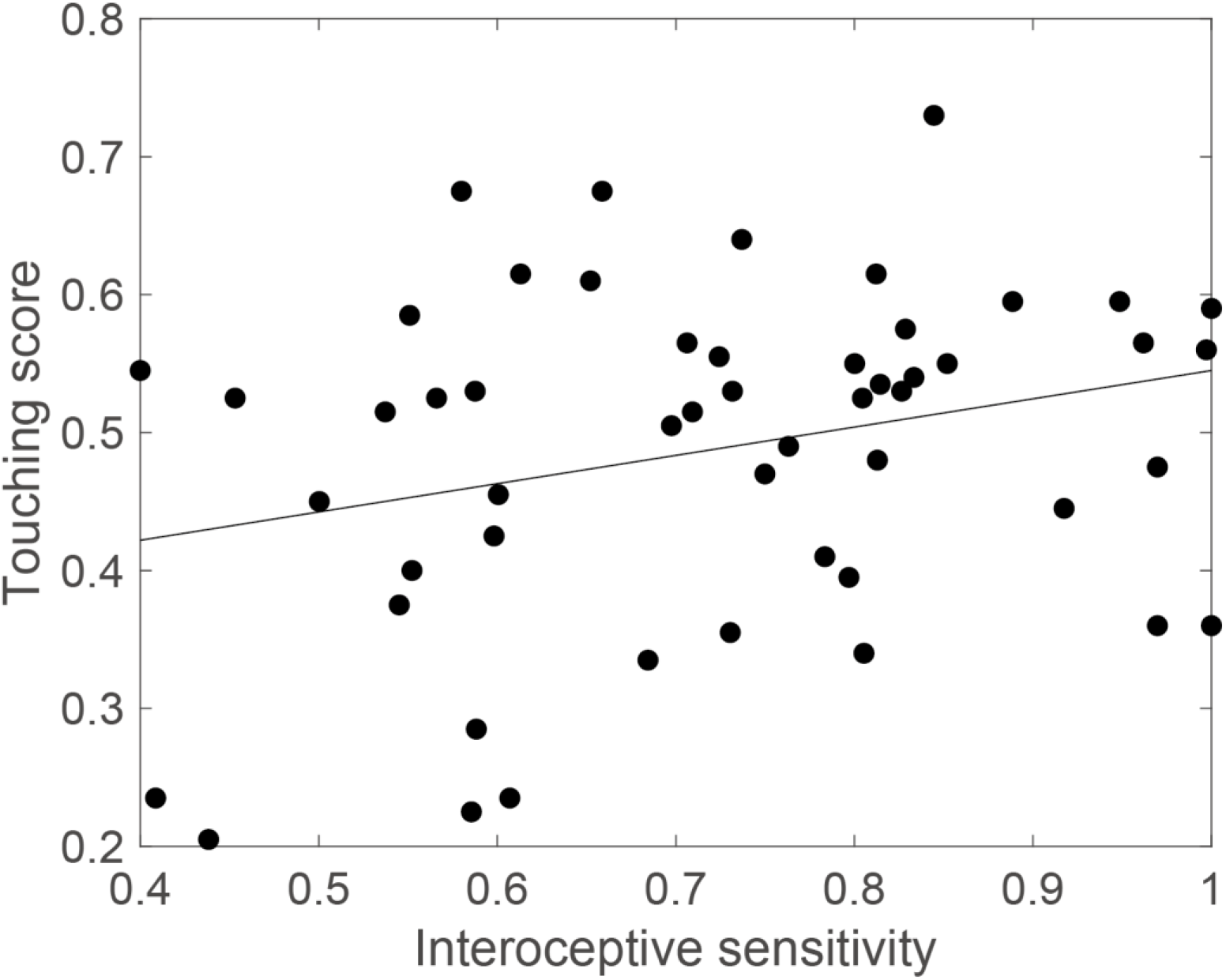
The relationship between the average touching score for all music pieces and interoceptive sensitivity defined by the amplitude parameter of the HDT. Each point indicates each participant’s data. The solid line shows a result of linear regression. HDT, heartbeat discrimination task.

### fMRI activity and interoceptive sensitivity

First, to investigate the activity related to the emotional experience induced by music, we compared whole-brain activity in the high-touching and low-touching trials. We observed significantly higher activation in the bilateral auditory cortices (superior temporal gyrus and inferior frontal gyrus), supplementary motor area, striatum (caudate nucleus, putamen, and globus pallidus), left primary motor area, left anterior insula, and left ventral inferior frontal cortex (Broca’s area) in the high-touching trials than in the low-touching trials (Figure 5, Table 2).

**Figure 5.**
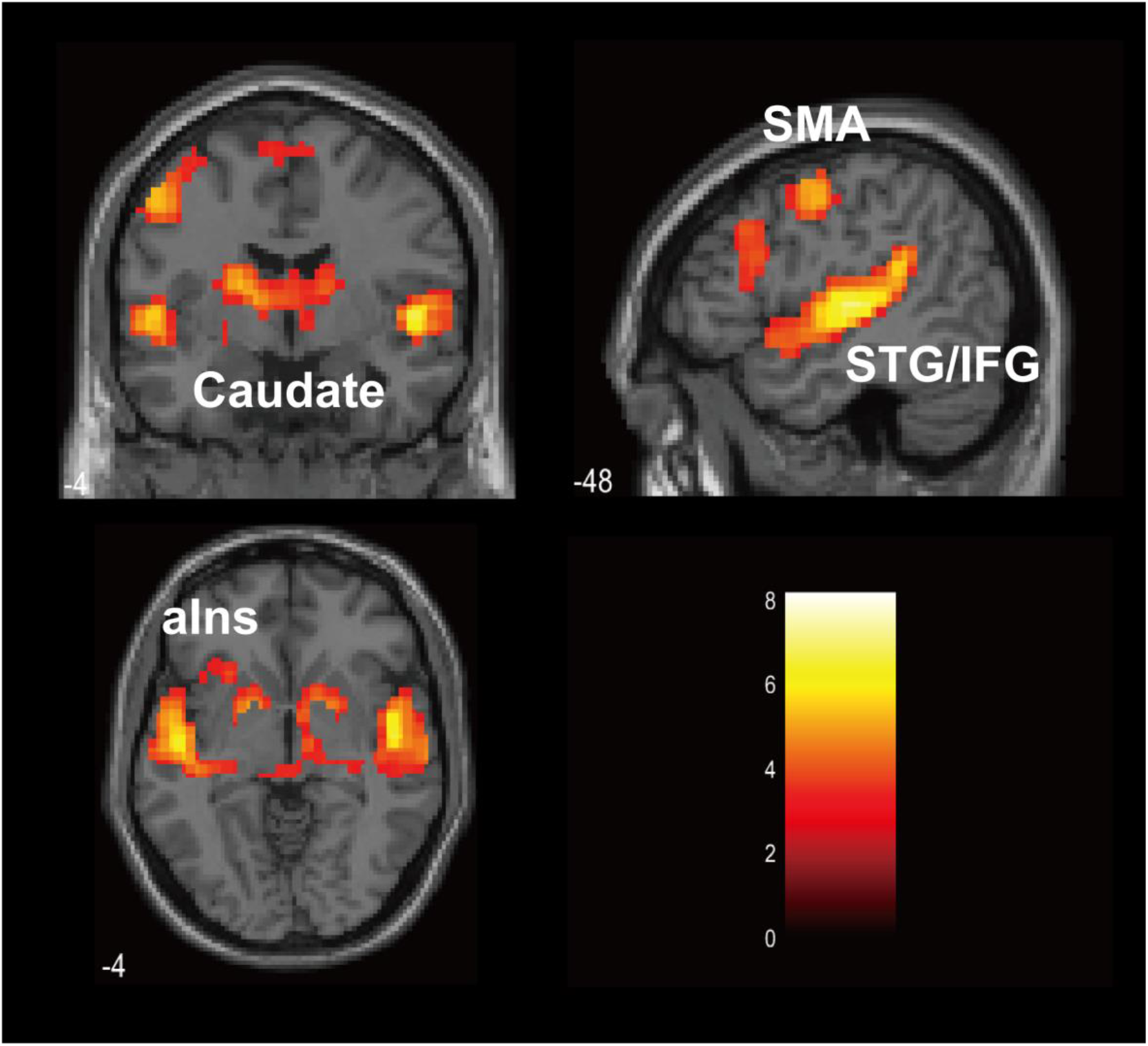
The brain regions that were more active when listening to music in the high-touching trials than in the low-touching trials (uncorrected p < 0.001 at voxel-level, FWE corrected p < 0.05 at cluster-level). SMA, Supplementary motor area; STG, Superior temporal gyrus; IFG, Inferior frontal gyrus; aIns, anterior Insula.

**Table 2.** Anatomical locations and coordinates of brain regions showing significant activations in the high-touching trials.

| Cluster | Region | Left hemisphere |  |  |  | Right hemisphere |  |  |  |
| --- | --- | --- | --- | --- | --- | --- | --- | --- | --- |
|  |  | t-score | x | y | z | t-score | x | y | z |
| Auditory cortex |  |  |  |  |  |  |  |  |  |
|  | Superior temporal gyrus | 7.42 | -51 | -16 | 4 | 8.17 | 51 | -13 | 0 |
|  | Inferior frontal gyrus | 5.07 | -42 | -31 | 13 | 4.08 | 54 | -22 | 13 |
| Primary motor cortex |  |  |  |  |  |  |  |  |  |
|  | Precentral gyrus | 5.33 | -51 | -1 | 48 |  |  |  |  |
| Supplementary motor area |  |  |  |  |  |  |  |  |  |
|  | Frontal superior gyrus | 5.20 | -9 | 17 | 48 | 3.92 | 3 | 5 | 64 |
| Broca's area / |  |  |  |  |  |  |  |  |  |
| Insular cortex |  |  |  |  |  |  |  |  |  |
|  | Frontal middle gyrus | 4.28 | -45 | 23 | 32 |  |  |  |  |
|  | Anterior insula cortex | 3.76 | -36 | 23 | -6 |  |  |  |  |
| Striatum |  |  |  |  |  |  |  |  |  |
|  | Caudate | 4.38 | -12 | 2 | 10 | 4.66 | 15 | 2 | 13 |
|  | Pallidum | 4.47 | -18 | 5 | 0 | 5.26 | 15 | 8 | -3 |
|  | Putamen | 4.54 | -18 | 8 | 0 | 5.11 | 21 | 5 | -3 |

The areas that were more active when listening to music in the high-touching trials than in the low-touching trials. Coordinates are derived from the standard brain of the Montreal Neurological Institute (MNI). The significance level is $p < 0.001$ without correction at voxel-level and $p < 0.05$ , with family wise error (FWE) correction at the cluster level.

Next, to investigate the relationship between individual differences in interoceptive sensitivity and activity in the insula subregions, we divided the participants into two groups, high-IS and low-IS groups, according to the heartbeat discrimination task parameter (*A*), as in the analysis of physiological responses. We conducted an ROI analysis of six insula subregions: hypergranular insula, ventral agranular insula, dorsal agranular insula, ventral dysgranular and granular insula, dorsal granular insula, and dorsal dysgranular insula, defined by the Human Brainnetome Atlas (Fan *et al.*, 2016; Table 3). For the ROI analysis, we used contrast images of high-touching and low-touching trials. We performed a *t*-test for the activities in the high-IS and low-IS groups. The *P*-value threshold was corrected with false discovery rate correction for multiple comparisons (*p* < 0.05). This analysis revealed that the activity in the dorsal granular insula in both hemispheres and ventral dysgranular and granular insula and dorsal dysgranular insula in the right hemisphere were significantly higher in the high-IS group than in the low-IS group (Figure 6, Table 3).

**Figure 6.**
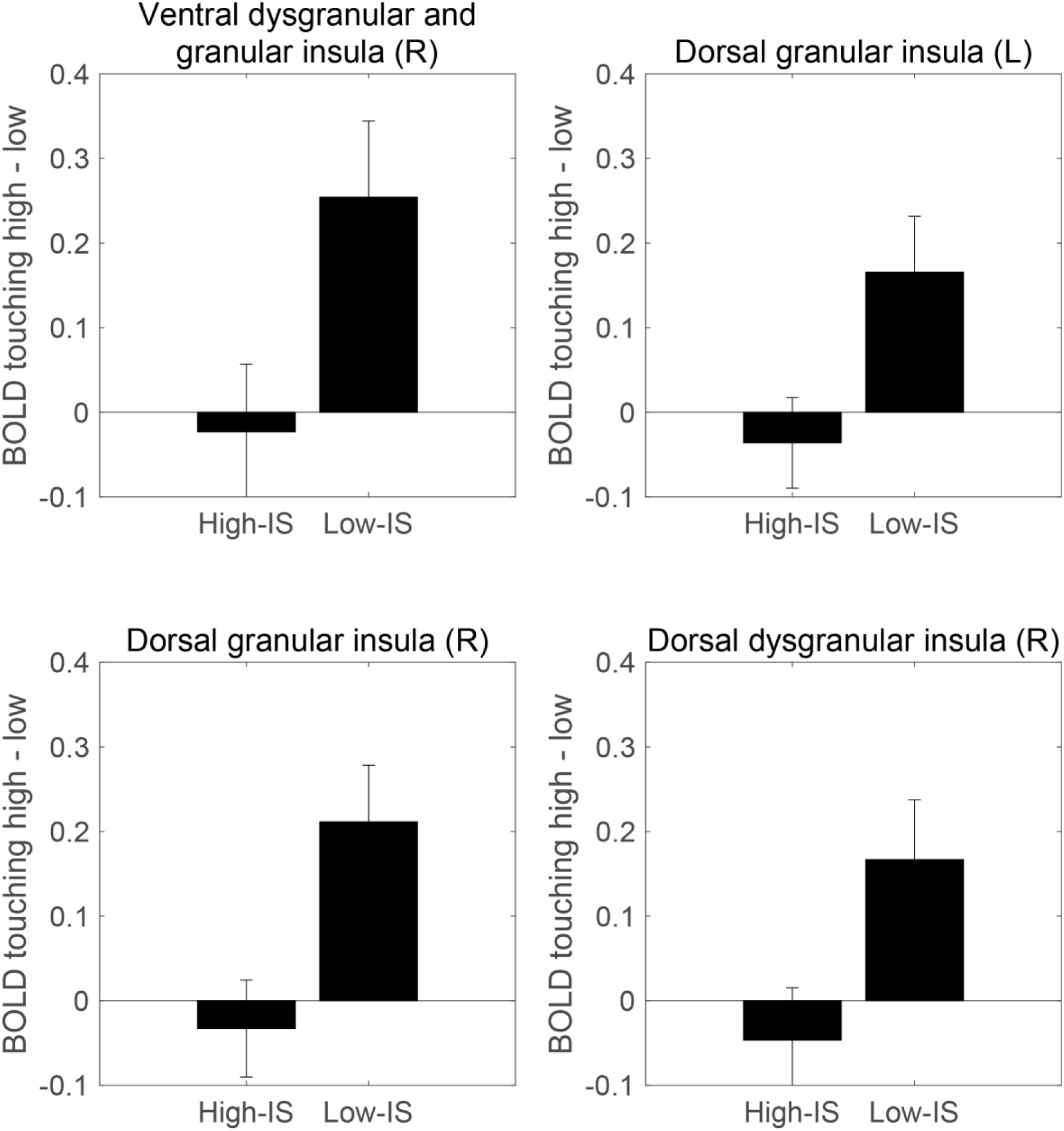
BOLD responses in the insula subregion for the low and high interoceptive sensitivity groups. Each figure represents the insula subregions showing significant differences between the high- and low-IS groups in the ROI analysis. The vertical axis represents the difference in the BOLD responses between the activity when listening to music in the high-touching trials and in the low-touching trials. The error bars represent ±1 standard error.

**Table 3.** Anatomical coordinates of insula subregions and t-value of ROI analysis.

|  | Left |  |  |  |  | Right |  |  |  |  |
| --- | --- | --- | --- | --- | --- | --- | --- | --- | --- | --- |
|  | t | sig. | x | y | z | t | sig. | x | y | z |
| hypergranular insula | 1.46 |  | -36 | -20 | 10 | 1.99 |  | 37 | -18 | 8 |
| ventral agranular insula | 0.56 |  | -32 | 14 | -13 | 1.85 |  | 33 | 14 | -13 |
| dorsal agranular insula | 1.38 |  | -34 | 18 | 1 | 1.24 |  | 36 | 18 | 1 |
| ventral dysgranular and granular insula | 1.18 |  | -38 | -4 | -9 | 2.31 | * | 39 | -2 | -9 |
| dorsal granular insula | 2.37 | * | -38 | -8 | 8 | 2.77 | * | 39 | -7 | 8 |
| dorsal dysgranular insula | 1.56 |  | -38 | 5 | 5 | 2.27 | * | 38 | 5 | 5 |

The regions of interest were defined using the Brainnetome Atlas. The $t$ -value indicates the result of a one-sided $t$ -test on whether the activity in the high-touching trials is larger in the high-IS group than in the low-IS group. The asterisk in the table shows $p < 0.05$ at the significance level corrected with false discovery rate correction for multiple comparisons. ROI,

## 4. Discussion

In this study, we assessed three hypotheses about emotional experience, physiological and brain activities, and the effect of individual differences in interoceptive sensitivity. (1) The physiological response during the emotional experience of music correlates with individual differences in interoceptive sensitivity. (2) The interoceptive sensitivity measured by the heartbeat discrimination task is related to the physiological response. (3) The activity of the posterior insula during emotional experience correlates with individual differences in interoceptive sensitivity. To estimate the participants’ interoceptive sensitivity, we used heartbeat discrimination and heartbeat counting tasks. Based on the newly created index (amplitude parameter, *A*) of the interoceptive sensitivity derived from the heartbeat discrimination task, the heart rate increased when listening to a high-touching music piece (high-touching trials) in the high interoceptive sensitivity group only (high-IS group). These findings support the first and second hypothesis that the physiological response during an emotional experience correlates with individual differences in interoceptive sensitivity, which was estimated by heartbeat discrimination task. The fMRI analysis showed that the left anterior insula was more active in the high-touching trials than in the low-touching trials. Furthermore, the ROI analysis of the insula subregions revealed that the high-IS individuals showed high activity in the mid-posterior insula in the high-touching trials. These results indicate that the posterior insula activity during an emotional experience correlates with the individual difference in the interoceptive sensitivity estimated by the heartbeat discrimination task, supporting the third hypothesis.

### Method for measuring interoceptive sensitivity

In this study, heartbeat counting and heartbeat discrimination tasks were performed by the same participants. In the comparison between the two tasks, a weak positive correlation (*r* = 0.27) was found between the IA and amplitude of the heartbeat discrimination task only. In previous studies, the correlation between heartbeat counting and heartbeat discrimination tasks has been controversial; some researchers have reported positive correlations (Garfinkel *et al.*, 2015; Garfinkel *et al.*, 2016; Kandasamy *et al.*, 2016; Rae *et al.*, 2019), whereas others have reported no correlation (Forkmann *et al.*, 2016; Ring and Brener, 2018; Herman *et al.*, 2019). An integrative review of these studies suggests a weak correlation of 0.20 (Hickman *et al.*, 2020), which is close to that of the present study. Contrary to the previous studies, we devised two new heartbeat discrimination indices (*A* and σ). The amplitude may represent the sensitivity to the heartbeat intensity, and the variance may reflect the goodness of the temporal resolution of heartbeat perception. Since the IA of the heartbeat counting task was correlated with the amplitude of the heartbeat discrimination task, our results suggest that the heartbeat counting task can estimate the sensitivity to heartbeat intensity.

Regarding the relationship between the music listening task and the heartbeat tasks, a significant difference in heart rate was observed only when the participants were grouped according to the heartbeat discrimination task amplitude. One reason for this finding is that the heartbeat discrimination task amplitude may have been more strongly related to the touching of the music. The heartbeat counting task requires good interoceptive sensitivity because it merely counts the heartbeat. In contrast, to perform the heartbeat discrimination task, participants must compare their heart beats with the sounds presented concurrently. Therefore, the heartbeat discrimination task requires integrated processing of interoception and exteroception. Notably, the generation of emotions requires the integration of interoception and exteroception (Barrett *et al.*, 2004; Barrett and Simmons, 2015); thus, the ability to integrate interoception and exteroception may be strongly associated with the physiological response to music.

### Individual differences in interoceptive sensitivity and physiological response

A comparison of the heart rates when listening to music among the high-touching, middle-touching, and low-touching trials showed that the higher the touching rating, the higher the heart rate. This is consistent with the typical tendency observed when listening to music or feeling emotions induced by music (Koelsch and Jäncke, 2015). Furthermore, regarding the relationship between changes in heart rate and individual differences in interoceptive sensitivity, it was only in the high-IS group that the heart rate was higher when listening to the high-touching music piece. This finding is consistent with those of previous studies comparing interoceptive sensitivity with the magnitude of the physiological response caused by visually induced emotions (Pollatos *et al.*, 2007; Pollatos and Schandry, 2008; Herbert *et al.*, 2010; De Witte *et al.*, 2016). However, no significant change in the heart rate was observed when we divided trials by the type of music pieces (tonal, atonal, and dissonance). This suggests that subjective evaluation is related to the heart rate. In addition, there was a weak positive correlation between interoceptive sensitivity and the average touching score, but no significant correlation between interoceptive sensitivity and the variance of touching score, the average heart rate, and the variance of heart rate was observed. This indicates that participants with low and high interoceptive sensitivity showed similar changes in touching scores and heart rate.

The finding that only the high-IS group had a relationship between the touching score and the heart rate has two possible interpretations. First, the heart rate of the high-IS individuals fluctuate greatly depending on the emotions. Second, the high-IS individuals refer to the heart rate when assessing emotions. If the first hypothesis was true, participants in the low-IS group would show less heart rate changes among the trials. However, no difference in the variance of heart rate changes between the high- and low-IS groups was observed. Thus, the second interpretation is more appropriate. Based on our findings, we can speculate that individuals with high interoceptive sensitivity use their physical condition to assess their emotional level when listening to music, while those with low interoceptive sensitivity use their knowledge of moving and non-moving music or experience of emotional music.

Dunn *et al.* (2010) showed that participants with high interoceptive sensitivity had a greater effect of anticipatory EDA on risk-aversion behavior. However, there was no difference in the magnitude of EDA depending on interoceptive sensitivity. In other words, when making a risky selection, EDA occurs in the same way regardless of the interoceptive sensitivity, but an individual with a high interoceptive sensitivity can effectively utilize it for selection behavior. Consistent with this result, in the present study, individuals in the high-IS group showed a large heart rate change during the high-touching trials, but the magnitude of heart rate change did not differ according to interoceptive sensitivity. Therefore, although the magnitude of the physiological response does not change depending on individual differences in interoceptive sensitivity, a difference in the use of the physiological response is apparent.

### Brain activity and interoceptive sensitivity

During the high-touching trials, we found significant activations in the bilateral auditory cortices, supplementary motor area, striatum, left primary motor area, anterior insula, and Broca’s area. The anterior insula and supplementary motor area are generally considered to be activated by emotional arousal (Lee *et al.*, 2015; Lindquist *et al.*, 2015). In addition, the brain activity related to music-induced emotions was observed in the striatum, amygdala, hypothalamus, cingulate cortex, insula, primary motor cortex, and prefrontal cortex (Koelsch, 2014). The activated areas in these previous studies are consistent with our findings, suggesting that the activity observed in our study was also caused by music-induced emotions. Furthermore, the activated areas involved in interoceptive processing overlap with those involved in emotion processing (Adolfi *et al.*, 2017; Quigley *et al.*, 2021), including the insula, inferior frontal gyrus, prefrontal cortex, striatum, supplementary motor area, and cingulate cortex. The activated areas in our study are consistent with these areas, which further supports the view that interoceptive activity in the brain is related to emotional activity.

Next, the ROI analysis of the insula subregions showed a significant difference between the high-IS group and low-IS group in the activity of the bilateral dorsal granular insula and the right ventral dysgranular and granular insula, as well as the right dorsal dysgranular insula. The anterior insula is thought to be responsible for cognitive and emotional functions, while the posterior insula is thought to be responsible for bodily sensorimotor functions, including internal organs. The anterior insula is connected to the orbitofrontal cortex, the cingulate cortex, and the limbic cortex, while the posterior insula is connected to the parietal area, such as the somatosensory and the motor cortices (Deen *et al.*, 2010; Cauda *et al.*, 2011; Cerliani *et al.*, 2012; Cloutman *et al.*, 2012), which correspond to their respective functions. Barrett and Simmons (2015) proposed a model of interoceptive processing focusing on the composition of the insula subregion and argued that the agranular cortex receives the interoceptive sensory input, while the granular cortex provides interoceptive predictions and is assigned an emotional function. Since the posterior insula, which showed a significant difference in activity between the high- and low-IS groups, receives sensory input, it is presumed that participants with high interoceptive sensitivity received more interoceptive sensory input during the emotional experience. From the result of physiological responses, we can speculate that participants with high interoceptive sensitivity received strong heartbeat sensory input and used it for emotional evaluation, and the strong sensory input may have been reflected in the activity of the posterior insula.

The mid-insula connects the anterior and posterior regions, and it has been suggested that the brain waves generated by pain are transmitted from the posterior insula to the anterior insula via the mid insula (Frot *et al.*, 2014). Kuehn *et al.* (2016) also showed that interoceptive sensitivity is related to the strength of the connection between the posterior insula and the mid/anterior insula. In this study, the strength of the connection between the anterior and posterior insula may be reflected in the activity of the mid insula.

## Conclusions

In this study, we investigated the relationship between individual differences in interoceptive sensitivity and ratings of emotional touch and heart rate when listening to music. We found increased heart rate in the high-touching trials for the high-IS group only, which was defined by the amplitude parameter of the heartbeat discrimination task. This finding suggests that individuals with high interoceptive sensitivity use their physical condition to assess their emotional level when listening to music. In addition, we found no significant change in the heart rate for cases where interoceptive sensitivity was defined by the heartbeat counting task, suggesting that the heartbeat discrimination task could be more strongly associated with the physiological response of emotional experience. Furthermore, our ROI analysis showed that the activities of the mid and posterior insula during the emotional experience were correlated with interoceptive sensitivity. This suggests that insula activity may reflect the use of interoception for estimating emotions. These results may extend the knowledge of interoceptive processing and elucidate the role of interoception in emotions.

## Acknowledgement

The authors are grateful to T. Tomita, H. Matsuura, and N. Miura for their assistance with data collection.

